# Lung epithelial cells have virus-specific and shared gene expression responses to infection by diverse respiratory viruses

**DOI:** 10.1101/090936

**Authors:** James T. VanLeuven, Benjamin J. Ridenhour, Craig R. Miller, Tanya A. Miura

## Abstract

The severity and outcome of respiratory viral infections is partially determined by the cellular response mounted by infected lung epithelial cells. Disease prevention and treatment is dependent on our understanding of the shared and unique responses elicited by diverse viruses, yet few studies compare host responses to different viruses while controlling other experimental parameters. We compared changes in gene expression of murine lung epithelial cells infected individually by three respiratory viruses causing mild (rhinovirus, RV1B), moderate (coronavirus, MHV-1), and severe (influenza A virus, PR8) disease in mice. RV1B infection caused numerous gene expression changes, but the differential effect peaked at 12 hours post-infection. PR8 altered an intermediate number of genes whose expression continued to change through 24 hours. MHV-1 had comparatively few effects on host gene expression. The viruses elicited highly overlapping responses in antiviral genes, though MHV-1 induced a lower type I interferon response than the other two viruses. Signature genes were identified for each virus and included host defense genes for PR8, tissue remodeling genes for RV1B, and transcription factors for MHV-1. Our comparative approach identified universal and specific transcriptional signatures of virus infection that can be used to discover mechanisms of pathogenesis in the respiratory tract.

## Introduction

Viruses from several different families, including *Picornaviridae, Orthomyxoviridae, Paramyxoviridae, Coronaviridae*, and *Adenoviridae*, infect and cause diseases in the respiratory tract. These diseases range from mild infections of the upper respiratory tract to severe diseases when infecting the lungs. Respiratory viruses commonly target epithelial cells of the airways and lungs. These epithelial cells are responsible for detecting viral pathogens and initiating antiviral responses at the level of infected cells and the immune system, and therefore their response to infection has an important role in determining disease outcomes. Knowledge of the shared and virus-specific responses of respiratory epithelial cells to infection by diverse viruses is fundamental to understanding viral pathogenesis and developing therapies to treat severe respiratory infections.

Murine models of respiratory viral infections have been widely used to identify the mechanisms that determine disease severity in the respiratory tract. While these models are invaluable for evaluating pathology and host responses to infection, parallel *in vitro* studies can be used to identify gene expression and signaling pathway changes that occur in infected cells to mediate pathogenesis. In this study, we compare the gene expression response of murine lung epithelial cells to infection by three respiratory viruses used in murine models: rhinovirus (serotype RV1B) in the family *Picornaviridae*, mouse hepatitis virus (MHV strain 1) in the family *Coronaviridae*, and influenza A virus (strain PR8) in the family *Orthomyxoviridae*. Viruses from different families and with different replication strategies were chosen to identify which responses to infection in respiratory epithelial cells are shared and which are virus-specific. In the following paragraphs, we describe some of the key features of these three viruses in murine models.

Minor group rhinoviruses, including RV1B, use low-density lipoprotein receptor to enter either human or murine cells. Because RV1B can replicate in mouse cells, it has been used to study immune responses and/or mechanisms of asthma exacerbation in infected mice [1-5]. Intranasal inoculation of mice with a high dose of RV1B results in viral replication and an early inflammatory response in the respiratory tract without clinical signs of disease [1, 3-5]. RV1B antigens have been detected in cells of the epithelia and lamina propria of the upper respiratory tract of infected mice [2, 4].

MHV-1 is a strain of mouse hepatitis virus that preferentially replicates and causes disease in the respiratory tract of specific mouse strains and thus serves as a model for respiratory coronaviral infections [6, 7]. MHV-1 causes severe disease in A/J-strain mice and milder pathology in other mouse strains [6, 8]. Mouse strain-dependent disease severity corresponds to inflammatory responses, fibrin deposition, and reduced type I interferon (IFN) production [6]. Further, type I IFN-mediated signaling is required for protection from severe disease during MHV-1 infection of resistant mice [8]. MHV-1 has been detected in pulmonary macrophages of infected mice, but has not been reported to infect respiratory epithelial cells *in vivo* [6]. Although primary murine alveolar epithelial cells are susceptible to MHV-1 infection *in vitro*, their role during *in vivo* infection is not clear [9].

Mice have been used for decades to study the pathogenesis of influenza viral disease. One of the most commonly used strains, PR8, has been serially passaged in mice to produce a model for pulmonary infection. PR8 infection results in a range of disease severities that is mouse strain-dependent [10]. Although susceptible mice mount a type I IFN response to PR8 infection, lethal infection is associated with spread of virus to the alveoli and an excessive inflammatory response [10-13]. PR8 replicates in bronchiolar and alveolar epithelial cells of the lower respiratory tract *in vivo* and in primary murine respiratory epithelial cells *in vitro* (Blazejewska et al., 2011; [9, 14, 15].

We used a murine lung epithelial cell line (LA4) to compare the gene expression response to these three unrelated viruses. LA4 cells were derived from neoplastic lung epithelia from strain A (A/He) mice and have some properties of alveolar type II cells [16]. Strain A (A/J) mice are highly susceptible to respiratory viral infections, including MHV-1 and influenza A viruses [6, 10]. Other studies have demonstrated that LA4 cells are susceptible to infection by PR8 and RV1B [15, 17]. In this study, we show that LA4 cells are also susceptible to infection by MHV-1 (hereafter referred to as MHV). The gene expression response of LA4 cells to infection by MHV, PR8, and RV1B (hereafter referred to as RV) differed in timing and magnitude of the changes. While we expected to see highly divergent transcription responses to these three viruses, they induced expression of a large overlapping set of shared genes, including genes involved in antiviral responses. Each virus also altered expression of unique genes, which highlight their different replication strategies and mechanisms of pathogenesis in murine hosts.

## 2. Materials and Methods

### 2.1. Virus stocks and cell lines

PR8 (A/Puerto Rico/8/1934 (H1N1)), obtained from BEI Resources (NR-3169), was grown and titrated by plaque assay in MDCK (ATCC: CCL-34) cells. MHV, obtained from ATCC (VR-261), was grown and titrated by plaque assay in 17Cl.1 cells [18] (provided by Dr. Kathryn Holmes, University of Colorado Denver School of Medicine). RV, obtained from ATCC (VR-1645), was grown and titrated by tissue culture infectious dose 50% (TCID_50_) assay in HeLa cells (ATCC: CCL-2). LA4 (ATCC: CCL-196), a murine lung epithelial cell line, was cultured in Ham's F12K medium (Mediatech, Manassas, VA).

### 2.2. Epithelial cell infection and microarray sample

Our experimental approach was to infect LA4 cells with the three viruses at times t=0 h and t=12 h and harvest RNA for microarray analysis at t=24 h. Controls were mock-inoculated at both time points. Preliminary experiments were done to establish a multiplicity of infection (MOI) for each virus that resulted in comparable numbers of cells positive for viral antigen at 24 h post-infection (Figure S1). Based on this, LA4 cells were inoculated with 3 TCID_50_/cell RV, 1 PFU/cell PR8, or 3 PFU/cell MHV. Triplicate wells of LA4 cells in 6-well plates were inoculated with each virus diluted in serum-free medium or were mock-inoculated with serum-free medium for 1 h at 37°C. Viral inocula were removed and the cells were rinsed twice with serum-free medium. The cells were incubated in Ham's F12K medium with 2% FBS for 12 or 24 h, at which time RNA was isolated from cell cultures using an RNAeasy Plus kit (QIAGEN) and transcript levels were measured by microarray (NimbleGen *Mus musculus* MM9 Expression Array). For the 24 h samples, the media were removed and replaced with fresh media 12 h after inoculation. Raw and processed data are available from NCBI Gene Expression Omnibus under accession number GSE89190.

### 2.3. Data analysis

NimbleScan v2.5 software (NibleGen, Madison, WI) was used to extract raw intensity data for each probe on each array. Intensity data were read into the R statistical computing environment and checked for quality [19]. Data were prepared for processing using the pdInfoBuilder package and then normalized using the robust multichip average (RMA) function in the oligo package [20].

Statistical tests for differences in expression between treatments were conducted on the normalized expression data using a linear mixed-effect model followed by linear contrasts corrected for multiple comparisons. More specifically, expression was modeled as a function of treatment while probes for a particular gene were treated as a random effects using the nlme::lme function in R. The data contained seven treatments: three viruses tested at two time points (12 h and 24 h) each plus the mock treatment (RV_12_, RV_24_, MHV_12_, MHV_24_, PR8_12_, PR8_24_, and mock). The following nine post hoc, two-sided contrasts were then performed on the fitted models using the multcomp::glht function in R: each virus-time combination vs. mock (RV_12_ vs. mock, RV_24_ vs. mock, MHV_12_ vs. mock, MHV_24_ vs. mock, PR8_12_ vs. mock, PR8_24_ vs. mock) and each pairwise combination of viruses at the 24 h time point (RV_24_ vs. MHV_24_, RV_24_ vs. PR8_24_ MHV_24_ vs. PR8_24_). These 9 tests have their *p*-values adjusted by the multcomp::summary.glht function according to their joint distribution. Any factors detected to be significant at the family (gene) level were then subsequently corrected using the Benjamini-Hochberg algorithm [21] with a false discovery rate set at 1%.

Genes associated with type I IFN responses were identified among the sets of genes with differential expression for each virus compared to mock at 12 h and 24 h using the Interferome v.2.01 database (http://interferome.its.monash.edu.au; [22]. This database was queried using the search criteria: Type I IFN (all), *in vitro, Mus musculus*, 2.0 fold change (up or down). Interferome genes were manually sorted into functional categories: antiviral, IFN signaling, viral detection, immune response, MHC class I, inhibitory, apoptosis, ubiquitination, miscellaneous, and unknown. The significance of each virus having genes in the specific categories was tested using a chi-squared test.

Gene expression responses to RV1B were compared between our data from mouse cells and published data using human cells [23] using the MGI vertebrate homology database provided by The Jackson Laboratory [24] as well as the annotate package in R.

## 3. Results and Discussion

### 3.1. MHV, PR8, andRValter cellular gene expression by different magnitudes and with different timing

In order to compare how unrelated respiratory viruses (MHV, PR8, and RV) alter gene expression of murine epithelial cells, we infected LA4 cells with each of the three viruses and evaluated cellular gene expression by microarray analysis at 12 and 24 h after infection compared to mock-inoculated controls. Figure 1 shows the log_2_-fold change in expression level of genes that were differentially expressed in virus-infected, compared to mock-inoculated, cells. By plotting the changes in gene expression at 12 vs. 24 hours, we observed differences in magnitude and timing of gene expression changes mediated by the three viruses. The genes with significantly different expression in MHV-infected cells had low fold change values (Figure 1A). At 24 h, when gene expression changes were the highest, genes that were up-regulated by MHV infection had log_2_-fold change values of less than five. In contrast, PR8 and RV induced expression of many genes by greater than five log_2_-fold at 24 h, and genes were spread consistently across the full range of values. By 24 h, the genes most strongly up-regulated by PR8 and RV induced changes of 7 - 9.5 log_2_-fold and 6 - 7.5 log_2_fold compared to mock, respectively. This same pattern was observed with the down-regulated genes (Figure 1).

**Figure 1.**
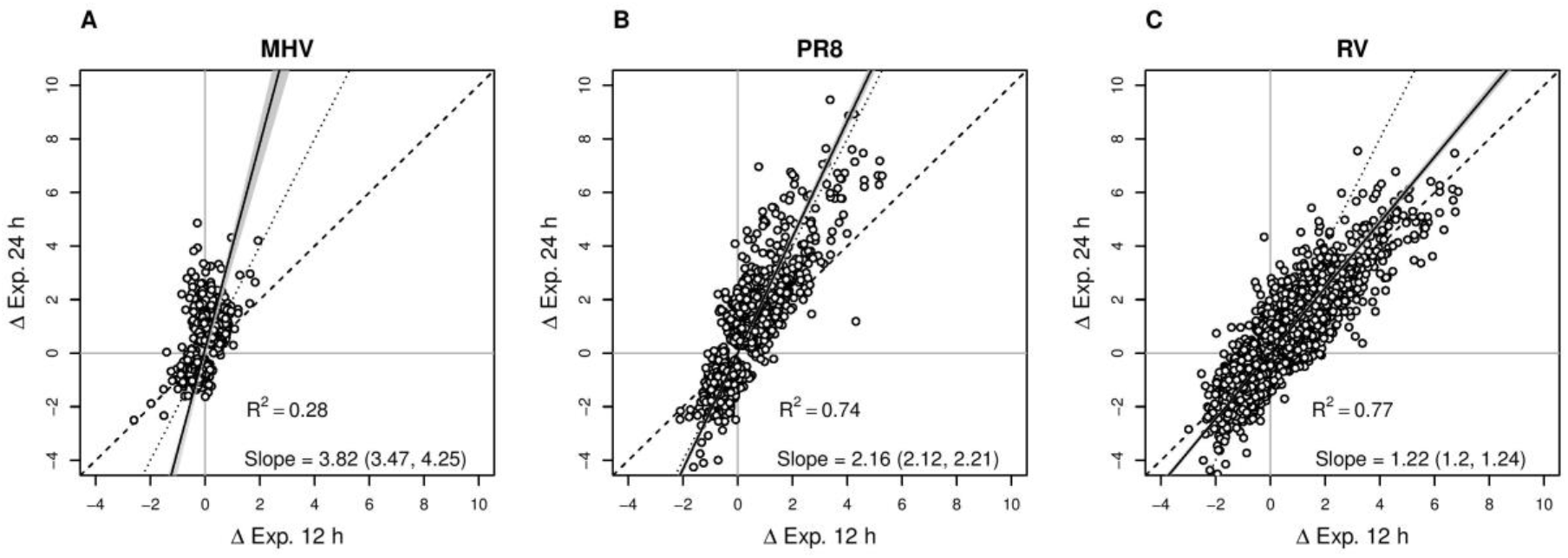
Magnitude and timing of gene expression changes mediated by MHV, PR8, and RV. The plots show the estimated log_2_ fold change in expression relative to mock at 12 h vs. 24 h for each of the three viruses. Each point represents one gene; only those genes that differ significantly from mock (at either time point) are included. The solid black line is the best fit regression line with the gray shading showing the 95% confidence interval. The slope with confidence interval and R^2^ are given in the inset legend. The dashed line illustrates the hypothesis that all changes in expression occur in the first 12 h (slope = 1); the dotted line shows the constant rate of change hypothesis (slope = 2). (a) MHV has small effects and most of the expression changes occur between 12 and 24 h (slope >> 2). (b) PR8 has larger effects than MHV and the changes approximate a constant rate of change across both 12 h intervals (slope ≈ 2). (c) RV also has large effects on gene expression and changes occur in the first 12 h with very little further change in the next 12 h (slope ≈ 1).

The three viruses also differed in the timing of gene expression changes. MHV altered expression of relatively few host genes, most of which were only significantly different from mock at 24 h (Figure 1A). While both PR8 and RV induced expression of large subsets of host genes, they did so with different timing. PR8-induced changes in gene expression occurred at a constant rate: the expression level of most genes at 24 h was approximately twice the expression value at 12 h (Figure 1B). In contrast, RV infection altered expression of a large number of genes by 12 h and the expression levels were maintained at approximately the same levels at the 24 h time point (Figure 1C).

Taken together, we observed differences in magnitude and timing of gene expression changes mediated by the three viruses: MHV changes were low and slow, PR8 induced gene expression to high levels at a steady rate, and RV altered gene expression more quickly to peak levels by 12 h. The limited response to MHV infection is in agreement with other coronaviruses, such as MHV-A59 [25] and SARS-CoV [26, 27]. In addition to inducing minor transcriptional up-regulation of host genes, MHV-A59 shuts down host gene expression by enhancing mRNA degradation [25]. A related coronavirus, SARS-CoV, also induces degradation of host mRNAs [28]. The low numbers of host mRNAs that were altered in response to MHV infection in our study could be due to one or both of these mechanisms. While rhinoviruses are also known to down-regulate host gene expression by inhibiting transcription, we saw a robust increase in host RNAs early upon RV infection. This is in agreement with other transcriptome studies of major and minor serogroup rhinoviruses in human respiratory epithelial cells and experimental infections of humans [23, 29-32]. The plateau in gene expression changes in RV-infected cells at 24 h may be due to transcriptional inhibition later in infection. PR8 infection induced a strong transcriptional response in LA4 cells, which has also been seen with multiple strains of influenza A viruses in primary human and mouse airway or lung epithelial cells [33-36].

### 3.2. Host genes have shared and unique responses to RV, PR8, and MHV infection

We identified which genes were altered by each virus at 24 h compared to mock and the degree of overlap among the differentially expressed genes. As was also observed in Figure 1, at 24 h RV infection resulted in up-regulation of the largest number of genes, followed by PR8 then MHV (Figure 2A). A similar pattern was seen with down-regulated genes (Figure 2B). While one might worry that the small number of significant genes that were altered by MHV could be false positives, the majority of these genes (65% of up-regulated and 86% of down-regulated genes) were also significantly altered by at least one other virus suggesting that most of these genes are true positives. For both up- and down-regulated gene sets, RV had the largest proportion of unique genes, while the majority of genes affected by both PR8 and MHV were shared by at least one other virus.

**Figure 2.**
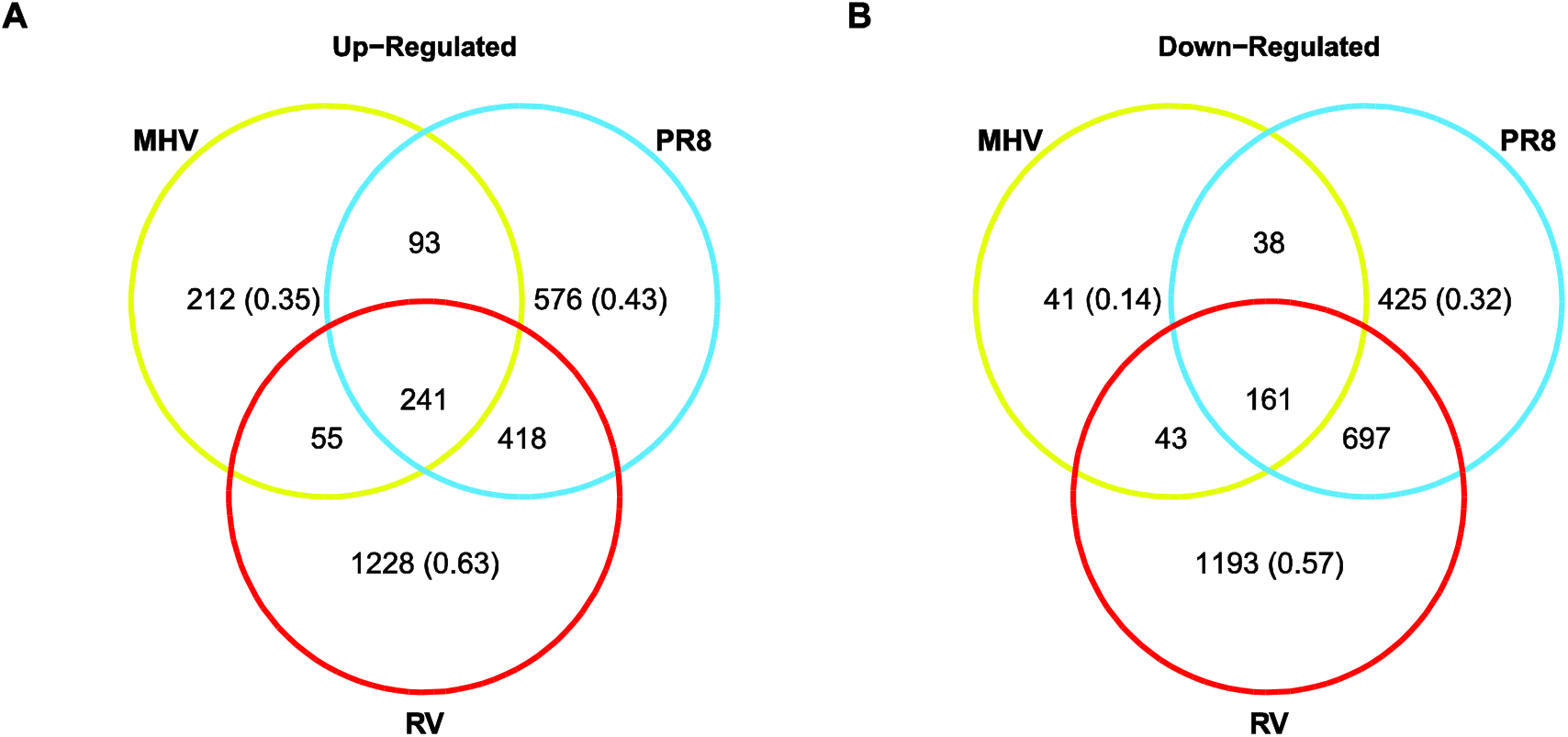
Numbers of genes with significantly altered expression upon viral infection. Venn diagrams show the number of significantly **(a)** up- and **(b)** down-regulated genes compared to mock 24 h after infection. The proportion of genes that are uniquely significant for each virus is indicated in parentheses.

Supplemental Table 1 contains the list of genes whose expression was significantly up-regulated by all three viruses compared to mock-inoculated cells. These genes may reflect a global response of epithelial cells to viral infection. Several of the genes with the highest fold change values are involved in antiviral defense at the level of infected cells (eg., *Mx1, Bst2, Oas2, Gbp10)* or recruitment of immune cells (eg., *Cxcl10, Cxclll*, *Cxcll).* These genes are upregulated by type I IFNs, suggesting that induction of a type I IFN response is shared by these viruses. In contrast to the shared up-regulated genes, genes that were significantly down-regulated by all three viruses have diverse functions (Table S2). Some examples of genes that were down-regulated by all three viruses included genes that encode transmembrane proteins *(Tmem 119, 231, 19, 50a*, and *14c)*, extracellular matrix proteins *(Spon2, Ogn, Aspn)*, and apoptotic signaling proteins *(Sdpr, Bmf, Bnip3l).*

### 3.3. Identification of signature genes that were uniquely altered by each virus

Comparing the number of genes altered by each virus provides insight into shared and unique cellular responses elicited by the viruses, but it does not provide information on the relative magnitudes of gene expression changes between viruses. To compare gene expression changes between viruses, we plotted the log_2_-fold change of each gene at 24 h for MHV vs. RV vs. PR8 (Figure 3A). We only included genes that were differentially expressed in at least one viral infection compared to mock. Like Figure 1, this 3D plot illustrates that PR8 and RV not only caused a larger number of genes to be up-regulated compared to MHV, but they also induced higher fold change values (Figure 3A).

**Figure 3.**
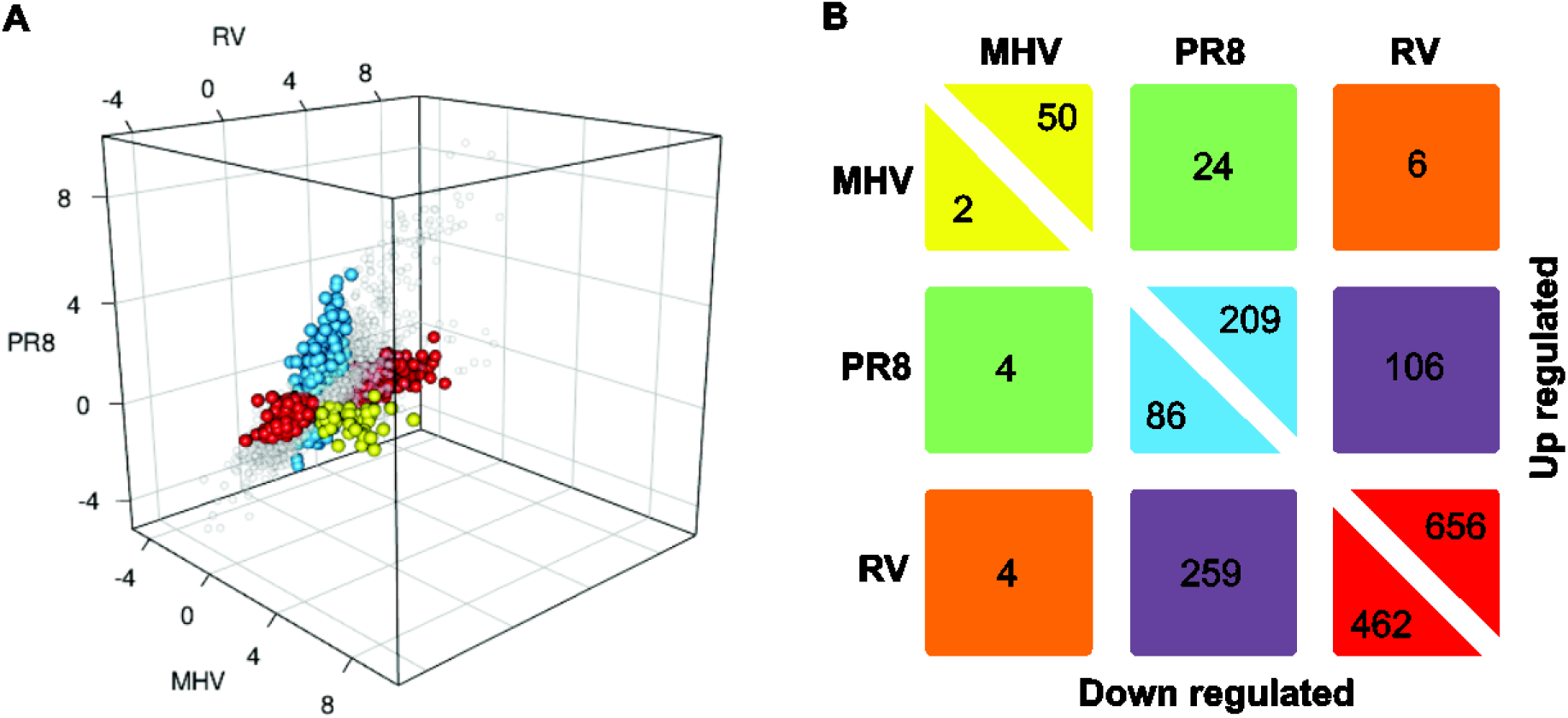
Patterns of gene expression changes mediated by viral infection. **(a)** Genes differentially expressed in at least one viral infection at 24 h are plotted as log_2_ fold change with each virus along a different axis. Signature genes, which have significantly larger effects in one virus compared to all other treatments, are colored: blue = PR8, red = RV, yellow = MHV. **(b)** The number of genes uniquely up- or down-regulated by each virus or pairs of viruses. The numbers along the center diagonal are the signature genes with boxes colored as in (a). The off-diagonal numbers are genes that have differential expression in two viruses compared to mock and the third virus, but are not significantly different from each other.

For each of the three viruses, we defined a signature gene as a gene that is both differentially regulated at 24 h compared to the mock treatment and has an effect size significantly larger than the other two viruses (i.e. fold change on the X axis is significantly different from Y-axis, Z-axis, and mock). These genes are colored in Figure 3A and appear along the diagonal in Figure 3B. As expected, RV had the largest number of signature genes, followed by PR8, then MHV (Figure 3B). Interestingly, the genes with the highest fold change values compared to mock were not signature genes, but were up-regulated by both PR8 and RV infection. A pairwise analysis was performed to identify the number of genes with altered expression compared with mock in two viruses compared with the third. This analysis, shown in Figure 3B, reveals that RV and PR8 had the most similarities in both up- and down-regulated genes (Figure 3B, purple blocks). The pattern of up-regulated gene expression changes during MHV infection was more similar to PR8 (24 genes) than RV (6 genes).

Several host defense genes were identified as signature genes uniquely up-regulated by PR8 infection (Table S3). These genes included cytokines and chemokines *(Cxcl9, Ccl5, IL12b, Ccl8)*, IFN response genes *(Ifitm6, Ifi27l2a, Ifna2, Ifit2, Ifitm5, Ifra11*), and genes involved in processing MHC class I antigens *(Psmb10, Tap2, H2-Q2, H2-K1, Psmd9, Psme2, Psme1).* The significant up-regulation of host defense genes in response to PR8 in the LA4 cell line corresponds with the expression profile of murine type II alveolar epithelial cells in response to PR8 infection in mice [37]. Furthermore, strong up-regulation of immune response-related genes upon PR8 infection of mice correlates with disease severity [11]. Several genes that were uniquely down-regulated by PR8 are involved in cellular metabolic pathways *(Cdo1, Aldh1a7, Acad11, Hsd17b4)* or intracellular transport *(Myl6b, Ift88, Anxa8).*

Although RV induced expression of several genes involved in host defense, these were largely shared by PR8 so were not identified as signature genes. The signature genes up-regulated by RV included kallikrein-1 and 10 kallikrein-1-related peptidases and additional proteins involved in tissue remodeling (Table S4). Rhinovirus infections are a significant cause of asthma exacerbations, which correspond with inflammatory responses in the airways. Kallikreins generate kinins and contribute to many disease processes, including inflammation. Kinins are induced by rhinovirus infections and kallikrein-1 is up-regulated by rhinovirus infection in humans, especially those with asthma [38, 39]. Up-regulation of these genes in mouse cells upon RV infection would provide a tractable animal model in which to study the roles of kallikreins in rhinovirus-induced asthma exacerbations. Rhinoviruses are also known to up-regulate expression of mucins by airway epithelial cells *in vitro* and *in vivo*, which may contribute to mucus hypersecretion [1, 40]. *Muc2* was the only mucin gene up-regulated by RV in our study, and was unique to RV infection (Table S4).

MHV infection resulted in regulation of a small set of signature genes (Figure 3B, Table S5). Signature genes that were uniquely up-regulated by MHV infection included multiple transcription factors from the double homeobox *(Duxf3, Dux, Dux4)* and zinc finger and SCAN domain *(Zscan4d, Zscan4c, Zscan4-ps1, 2 and 3)* families. Despite up-regulating expression of transcription factors, MHV infection had a minor impact on the host cell transcriptome. This may be due to enhanced degradation of mRNAs as discussed above, which has been shown to occur during other coronaviral infections [25, 28]. Therefore, LA4 cells may be up-regulating transcription in response to MHV infection through expression of various transcription factors while MHV causes degradation of these transcripts, which would reflect the small number of up-regulated transcripts in MHV infected samples. In contrast to MHV-A59, MHV-1 infection did not cause down-regulation of a substantial number of host genes. Differences may be due to virus strain, host cell type, and timing differences between the studies.

### 3.4. Type IIFN-relatedgenes had increased expression in LA4 cells infected by PR8, RV, and MHV

As described above, several of the genes with up-regulated expression in response to all three viruses and those that were unique to PR8 are induced by type I IFNs. To specifically evaluate how IFN response genes were altered by the three viruses, genes that were significantly up-regulated by each virus at the 24 h time point were used to query the Interferome v2.01 database (see Materials and Methods). A Venn diagram was generated to visualize the degree of overlap in IFN-related genes whose expression was induced by at least one of the three viruses (Figure 4). PR8 induced expression of the greatest number of IFN-related genes, a majority of which were shared by at least one other virus. RV up-regulated slightly fewer IFN-induced genes compared to PR8 and MHV infection resulted in up-regulation of the fewest IFN-induced genes. It was somewhat surprising that PR8 induced a higher type I IFN response than RV, given that RV induced expression of nearly twice as many genes than PR8 (Figure 2).

**Figure 4.**
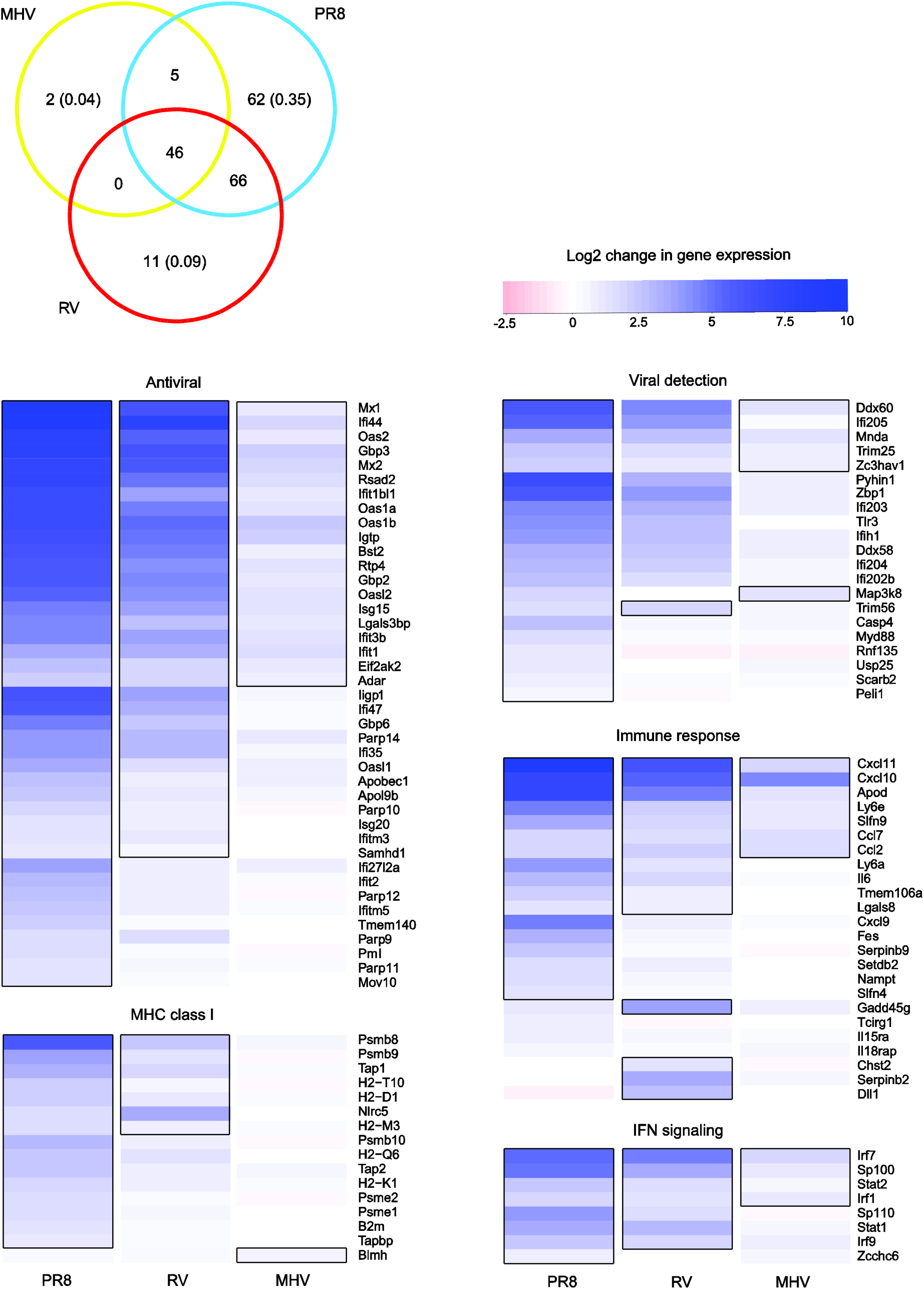
Differential expression of type I interferon-induced genes. Genes with significantly up-regulated expression compared to mock at 24 h (see Figure 2) were used to query the Interferome v.2.01 database. The Venn diagram shows the number of shared and unique type I IFN-related genes that were up-regulated in each viral infection. The proportion of genes up-regulated in only one virus treatment are shown in parentheses. The genes represented in the Venn diagram were divided into functional groups and heat maps were generated using log_2_ fold change values for each virus at 24 h compared to mock-inoculated controls. Heat maps of additional functional groups can be found in Supplemental Figure 2. Gene names are indicated to the right of each row and statistically significant values are outlined in black.

There was strong overlap between the IFN-induced genes up-regulated by each virus. The timing of IFN-related gene expression followed the same trend as was seen in Figure 1, wherein all genes with significantly altered expression were analyzed (data not shown). Most of the IFN-related genes up-regulated by MHV were only increased at 24 h. PR8 induced expression of 110 IFN-related genes at 12 h and these genes were a subset of the 179 genes up-regulated at 24 h. In contrast, RV infection induced expression of more IFN-related genes at 12 h (148 genes) than at 24 h (123 genes). Relative to up-regulation, few IFN-related genes were down-regulated at the 24 h time point (MHV=5, PR8=10, RV=26).

Type I IFNs induce expression of genes with different functions during an antiviral response. To determine whether there were specific patterns in expression of IFN-induced genes that correspond with function, the IFN-induced genes that had significantly increased expression by any of the three viruses were separated into functional groups. Heatmaps that demonstrate differences in fold change (color scale) and significant differences (outlined boxes) in expression compared to mock-inoculated controls were generated (Figure 4 and S2). As shown in the Venn diagram, this analysis also demonstrates that PR8 infection resulted in up-regulation of the most genes involved in type I IFN responses, followed by RV then MHV. The fold change values induced by PR8 infection also were generally higher than the other two viruses. However, there was not a significant correlation between virus identity and functional group. For most of the functional groups, MHV up-regulated expression of a smaller subset of the same genes as PR8 and RV, with the exception of the MHC class I pathway (Figure 4). MHV significantly up-regulated expression of only one gene involved in the MHC class I pathway *(Blmh)*, which was not significantly up-regulated by the other two viruses. This observation suggests that cytotoxic T cell responses may differ in MHV infections compared to PR8 and RV. T cell responses have complex roles in MHV-1 infections, as they contribute to protection in resistant mouse strains but mediate pathology in susceptible strains [41]. However, mice with the CD8+ T cell repertoire of a resistant strain in the background of a susceptible strain remain susceptible to severe MHV-1 infection [42]. The failure of MHV-1 to activate processing and presentation of MHC class I antigens could explain the inability of a broadly reactive CD8+ T cell response to protect these mice.

The interferome analysis focuses on IFN-induced gene expression, but not expression of the type I IFNs that induce these responses. Multiple type I IFNs exist, including IFN-p and 14 subtypes of IFN-a, all of which signal through the type I IFN a/p receptor (IFNAR) [43]. Type I IFNs can induce autocrine and paracrine signaling; thus the IFN-induced genes we detected could be from both infected and uninfected cells in the cultures. To determine if differential expression of type I IFNs explains the differences in IFN-induced gene expression upon infection by PR8, RV, and MHV, we analyzed the expression of type I IFN and receptor genes for each virus compared to mock (Figure 5). Probes for IFN-p 1 and ten subtypes of IFN-a were present on the arrays. In agreement with expression of IFN-induced genes, PR8 induced expression of the largest set of type I IFNs, followed by RV. Both viruses induced expression of *Ifnb* and *Ifoa4*, which encode type I IFNs known to be expressed early during antiviral responses [44, 45]. Five subtypes of *Ifna* were up-regulated by both PR8 and RV, while three *Ifna* subtypes were uniquely up-regulated by PR8 and only *Ifnab* was uniquely up-regulated by RV. Only PR8 induced expression of *Ifnar2*, which encodes the high affinity chain of the type I IFN a/p receptor [46]. This might allow for enhanced positive-feedback signaling and account for the larger number of IFN-induced genes up-regulated by PR8 infection.

**Figure 5.**
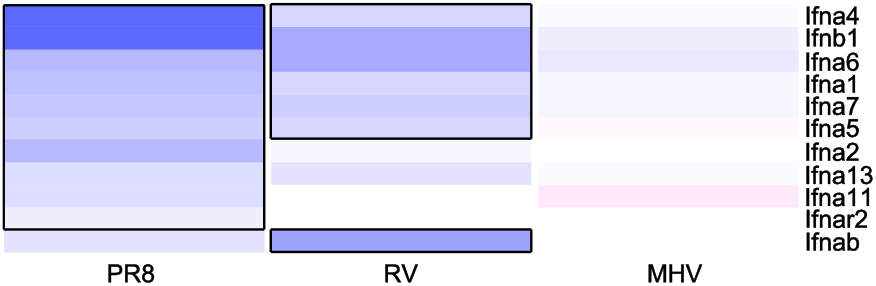
Differential expression of type I interferons and receptors. The log_2_ fold change compared to mock of up-regulated type I IFN cytokine and receptor genes. Gene names are shown to the right of each row and the color scale is the same as in Figure 4.

Rhinoviruses and influenza A viruses are known to induce type I IFN responses through recognition by MDA-5 and RIG-I, respectively [47, 48]. Furthermore, both viruses are recognized by TLR3 in infected epithelial cells [47, 48]. However, TLR3 predominantly induces expression of pro-inflammatory genes, rather than type I IFN-dependent genes, during influenza A virus infection [47]. Differential signaling through MDA-5 and RIG-I pathways may contribute to the differences in type I IFN responses by these two viruses. Zaritsky et al. have demonstrated that the type I IFN response to Sendai virus differs when cells are infected by different doses [49]. They further showed that these differences were mediated by differential signaling through the IFN a/p receptor, with robust signaling in uninfected cells. This supports our findings that PR8 induces expression of *Ifnar2* and additional type I IFN genes that are not up-regulated by RV (Figure 5).

None of the type I IFNs or receptors had significantly altered expression upon MHV infection (Figure 5), despite up-regulation of a modest number of IFN-stimulated genes (Figure 4). This could be due to IFN-independent expression of these genes, or induction by a type I IFN that was not represented on the microarray. Coronaviruses are notorious for being able to replicate within cells without triggering type I IFN responses, or delaying IFN induction until late in the replication cycle [34, 50-52]. Other studies have shown that the IFN response to MHV-1 is a critical determinant of susceptibility. Severe disease in A/J mice compared to C57Bl/6 mice correlates with lower type I IFNs detected in the lungs of A/J mice upon MHV-1 infection [6, 53]. Similarly, the expression of various type I IFNs in response to MHV-1 infection *in vitro* is cell line-dependent [53]. Because the cell line we used, LA4, was derived from the lungs of A/He mice, we would expect it to have a similar response as A/J mice. Thus the lack of type I IFNs induced by MHV-1 in LA4 cells *in vitro* corresponds with pathogenesis observed in A/J mice *in vivo.*

The finding that LA4 cells mount a stronger response to PR8 than RV or MHV infection may be due to differences in the viral recognition and signaling pathways used to detect these different viruses and amplification of the type I IFN response as discussed above. Alternatively, it could be due to differences in replication kinetics of the viruses. RV-infected cells have started dying by the 24 h time point (not shown), therefore host response genes may have been up-regulated at an earlier time point. In contrast, coronaviruses are known to delay cellular responses to infection [54] so the 24 h time point may be too early to evaluate the innate response to MHV infection. Alternatively, the cells may detect MHV and up-regulate transcription of IFN response genes, but mRNA degradation would mask this process. By quantifying mRNA transcripts at two time points after viral infection, our study cannot distinguish between these possibilities.

### 3.5. RV1B induced a similar gene expression response in murine and human respiratory epithelial cells

One limitation of our study is the analysis of three viruses that do not share a natural host. MHV is a natural pathogen of mice and PR8 is a highly mouse-adapted strain of influenza A virus. However, RV1B is a human rhinovirus whose receptor is conserved between mice and humans. RV1B is increasingly being used in mouse studies [1-4, 55]. Despite the difference in host, we found similar changes to gene expression in murine cells as studies with RV1B in human cells [23]. Of the 24,204 and 12,438 genes represented on our mouse microarray and the human microarray chip used by Chen et al., respectively, 10,847 genes are shared. Using the same 2-fold increase in expression cut-off and restricting our list only to homologous human genes studied by Chen et al., we found that 196 mouse genes were upregulated by RV1B infection. Comparing this list of 196 genes to the 48 upregulated human genes identified by Chen et al., we found that 20 genes (Table S6) were upregulated by RV1B infection in both human and mouse cells. A chi-squared test confirmed the significance of this shared pattern of up-regulated genes (χ^2^=431.7, d.f.=1, p<0.001). Interestingly, all 20 of the shared genes we identified are involved in type I IFN responses. While far from identical, the similarity of the responses in the two cell types suggests conserved activation of type I IFN responses by these different hosts and supports the validity of a murine model for studying rhinovirus infections in humans.

## 4. Conclusions

Alveolar epithelial cells have a key role in alerting the immune system to infection by respiratory viruses and shaping immune responses [37, 56, 57]. As viruses from several different families all target respiratory epithelial cells, it is important to understand the similarities and differences in how these cells respond to a diverse set of viruses. A significant number of genes were up-or down-regulated in response to infection by three distinct viruses from different families. Genes that were associated with a shared response to the three viruses included those involved in defense against viruses, and particularly genes that are induced by type I IFNs. However, there were differences in the timing, numbers of genes altered, and expression levels of these genes. This may reflect differences in viral replication cycles and signaling pathways that are activated by infection.

## Supplementary Materials

Figure S1: Infection of LA4 cells visualized by immunofluorescent assay of viral proteins and epifluorescent microscopy, Figure S2: Heatmap of differential expression values for interferome genes, Table S1: Genes up-regulated in all virus infections, Table S2: Genes down-regulated in all virus infections, Table S3: PR8 signature genes, Table S4: RV signature genes, Table S5: MHV signature genes, Table S6: Up-regulated genes shared with Chen et al., 2006.

## Acknowledgments

The authors are grateful to Dr. Kathryn Holmes, University of Colorado at Denver School of Medicine, Dr. Elizabeth Fortunato, University of Idaho, and Dr. Julian Leibowitz, Texas A&M University for cells and antibodies that were used in this study. The following reagents were obtained through the NIH Biodefense and Emerging Infections Research Resources Repository, NIAID, NIH: Influenza A Virus, A/Puerto Rico/8/34 (H1N1), NR-3169 and Polyclonal Anti-Influenza Virus H1 (H0) Hemagglutinin (HA), A/Puerto Rico/8/34 (H1H1), (Antiserum, Goat), NR-3148. Dr. Matthew Settles, Dr. Sam Hunter and Mr. Dan New in the IBEST Genomics Resources Core provided support with microarray processing and analysis. Ms. Ann Norton in the IBEST Optical Imaging Core provided support with microscopy.

## Author Contributions

T.M. and C.M. conceived and designed the experiments; T.M. performed the experiments; T.M., C.M., B.R., and J.V. analyzed the data; T.M., C.M., B.R., and J.V. wrote the paper.

## References

1. Bartlett, N. W.; Walton, R. P.; Edwards, M. R.; Aniscenko, J.; Caramori, G.; Zhu, J.; Glanville, N.; Choy, K. J.; Jourdan, P.; Burnet, J.; Tuthill, T. J.; Pedrick, M. S.; Hurle, M. J.; Plumpton, C.; Sharp, N. A.; Bussell, J. N.; Swallow, D. M.; Schwarze, J.; Guy, B.; Almond, J. W.; Jeffery, P. K.; Lloyd, C. M.; Papi, A.; Killington, R. A.; Rowlands, D. J.; Blair, E. D.; Clarke, N. J.; Johnston, S. L., Mouse models of rhinovirus-induced disease and exacerbation of allergic airway inflammation. Nat Med 2008, 14, (2), 199–204.

2. Lee, S. B.; Song, J. A.; Choi, G. E.; Kim, H. S.; Jang, Y. J., Rhinovirus infection in murine chronic allergic rhinosinusitis model. Int Forum Allergy Rhinol 2016.

3. Nagarkar, D. R.; Wang, Q.; Shim, J.; Zhao, Y.; Tsai, W. C.; Lukacs, N. W.; Sajjan, U.; Hershenson, M. B., CXCR2 is required for neutrophilic airway inflammation and hyperresponsiveness in a mouse model of human rhinovirus infection. J Immunol 2009, 183, (10), 6698–707.

4. Newcomb, D. C.; Sajjan, U. S.; Nagarkar, D. R.; Wang, Q.; Nanua, S.; Zhou, Y.; McHenry, C. L.; Hennrick, K. T.; Tsai, W. C.; Bentley, J. K.; Lukacs, N. W.; Johnston, S. L.; Hershenson, M. B., Human rhinovirus 1B exposure induces phosphatidylinositol 3-kinase-dependent airway inflammation in mice. Am JRespir Crit Care Med 2008, 177, (10), 1111–21.

5. Wang, Q.; Miller, D. J.; Bowman, E. R.; Nagarkar, D. R.; Schneider, D.; Zhao, Y.; Linn, M. J.; Goldsmith, A. M.; Bentley, J. K.; Sajjan, U. S.; Hershenson, M. B., MDA5 and TLR3 initiate pro-inflammatory signaling pathways leading to rhinovirus-induced airways inflammation and hyperresponsiveness. PLoS Pathog 2011, 7, (5), e1002070.

6. De Albuquerque, N.; Baig, E.; Ma, X.; Zhang, J.; He, W.; Rowe, A.; Habal, M.; Liu, M.; Shalev, I.; Downey, G. P.; Gorczynski, R.; Butany, J.; Leibowitz, J.; Weiss, S. R.; McGilvray, I. D.; Phillips, M. J.; Fish, E. N.; Levy, G. A., Murine hepatitis virus strain 1 produces a clinically relevant model of severe acute respiratory syndrome in A/J mice. J Virol 2006, 80, (21), 10382–94.

7. Leibowitz, J. L.; Srinivasa, R.; Williamson, S. T.; Chua, M. M.; Liu, M.; Wu, S.; Kang, H.; Ma, X. Z.; Zhang, J.; Shalev, I.; Smith, R.; Phillips, M. J.; Levy, G. A.; Weiss, S. R., Genetic determinants of mouse hepatitis virus strain 1 pneumovirulence. J Virol 2010, 84, (18), 9278–91.

8. Khanolkar, A.; Hartwig, S. M.; Haag, B. A.; Meyerholz, D. K.; Harty, J. T.; Varga, S. M., Toll-like receptor 4 deficiency increases disease and mortality after mouse hepatitis virus type 1 infection of susceptible C3H mice. J Virol 2009, 83, (17), 8946–56.

9. Kebaabetswe, L. P.; Haick, A. K.; Miura, T. A., Differentiated phenotypes of primary murine alveolar epithelial cells and their susceptibility to infection by respiratory viruses. Virus Res 2013, 175, (2), 110–9.

10. Srivastava, B.; Blazejewska, P.; Hessmann, M.; Bruder, D.; Geffers, R.; Mauel, S.; Gruber, A. D.; Schughart, K., Host genetic background strongly influences the response to influenza a virus infections. PLoS One 2009, 4, (3), e4857.

11. Alberts, R.; Srivastava, B.; Wu, H.; Viegas, N.; Geffers, R.; Klawonn, F.; Novoselova, N.; do Valle, T. Z.; Panthier, J. J.; Schughart, K., Gene expression changes in the host response between resistant and susceptible inbred mouse strains after influenza A infection. Microbes and infection /Institut Pasteur 2010, 12, (4), 309–18.

12. Blazejewska, P.; Koscinski, L.; Viegas, N.; Anhlan, D.; Ludwig, S.; Schughart, K., Pathogenicity of different PR8 influenza A virus variants in mice is determined by both viral and host factors. Virology 2011, 412, (1), 36–45.

13. Fukushi, M.; Ito, T.; Oka, T.; Kitazawa, T.; Miyoshi-Akiyama, T.; Kirikae, T.; Yamashita, M.; Kudo, K., Serial histopathological examination of the lungs of mice infected with influenza A virus PR8 strain. PLoS One 2011, 6, (6), e21207.

14. Kebaabetswe, L. P.; Haick, A. K.; Gritsenko, M. A.; Fillmore, T. L.; Chu, R. K.; Purvine, S. O.; Webb-Robertson, B. J.; Matzke, M. M.; Smith, R. D.; Waters, K. M.; Metz, T. O.; Miura, T. A., Proteomic analysis reveals down-regulation of surfactant protein B in murine type II pneumocytes infected with influenza A virus. Virology 2015, 483, 96–107.

15. Tate, M. D.; Schilter, H. C.; Brooks, A. G.; Reading, P. C., Responses of mouse airway epithelial cells and alveolar macrophages to virulent and avirulent strains of influenza A virus. Viral immunology 2011, 24, (2), 77–88.

16. Stoner, G. D.; Hallman, M.; Troxell, M. C., Lecithin biosynthesis in a clonal line of lung adenoma cells with type II alveolar cell properties. Experimental and molecular pathology 1978, 29, (1), 102–14.

17. Tuthill, T. J.; Papadopoulos, N. G.; Jourdan, P.; Challinor, L. J.; Sharp, N. A.; Plumpton, C.; Shah, K.; Barnard, S.; Dash, L.; Burnet, J.; Killington, R. A.; Rowlands, D. J.; Clarke, N. J.; Blair, E. D.; Johnston, S. L., Mouse respiratory epithelial cells support efficient replication of human rhinovirus. J Gen Virol 2003, 84, (Pt 10), 2829–36.

18. Sturman, L. S.; Takemoto, K. K., Enhanced growth of a murine coronavirus in transformed mouse cells. Infection and immunity 1972, 6, (4), 501–7.

19. Rosenblum, E. B.; Poorten, T. J.; Joneson, S.; Settles, M., Substrate-specific gene expression in Batrachochytrium dendrobatidis, the chytrid pathogen of amphibians. PLoS ONE 2012, 7, (11), e49924.

20. Carvalho, B. S.; Irizarry, R. A., A framework for oligonucleotide microarray preprocessing. Bioinformatics 2010, 26, (19), 2363–7.

21. Benjamini, Y.; Hochberg, Y., Controlling the False Discovery Rate - a Practical and Powerful Approach to Multiple Testing. J Roy Stat Soc B Met 1995, 57, (1), 289–300.

22. Rusinova, I.; Forster, S.; Yu, S.; Kannan, A.; Masse, M.; Cumming, H.; Chapman, R.; Hertzog, P. J., Interferome v2.0: an updated database of annotated interferon-regulated genes. Nucleic acids research 2013, 41, (Database issue), D1040–6.

23. Chen, Y.; Hamati, E.; Lee, P. K.; Lee, W. M.; Wachi, S.; Schnurr, D.; Yagi, S.; Dolganov, G.; Boushey, H.; Avila, P.; Wu, R., Rhinovirus induces airway epithelial gene expression through double-stranded RNA and IFN-dependent pathways. Am J Respir Cell Mol Biol 2006, 34, (2), 192–203.

24. Eppig, J. T.; Blake, J. A.; Bult, C. J.; Kadin, J. A.; Richardson, J. E.; Mouse Genome Database, G., The Mouse Genome Database (MGD): facilitating mouse as a model for human biology and disease. Nucleic acids research 2015, 43, (Database issue), D726–36.

25. Raaben, M.; Groot Koerkamp, M. J.; Rottier, P. J.; de Haan, C. A., Mouse hepatitis coronavirus replication induces host translational shutoff and mRNA decay, with concomitant formation of stress granules and processing bodies. Cellular microbiology 2007, 9, (9), 2218–29.

26. Josset, L.; Menachery, V. D.; Gralinski, L. E.; Agnihothram, S.; Sova, P.; Carter, V. S.; Yount, B. L.; Graham, R. L.; Baric, R. S.; Katze, M. G., Cell host response to infection with novel human coronavirus EMC predicts potential antivirals and important differences with SARS coronavirus. MBio 2013, 4, (3), e00165–13.

27. Sims, A. C.; Tilton, S. C.; Menachery, V. D.; Gralinski, L. E.; Schafer, A.; Matzke, M. M.; Webb-Robertson, B. J.; Chang, J.; Luna, M. L.; Long, C. E.; Shukla, A. K.; Bankhead, A. R., 3rd; Burkett, S. E.; Zornetzer, G.; Tseng, C. T.; Metz, T. O.; Pickles, R.; McWeeney, S.; Smith, R. D.; Katze, M. G.; Waters, K. M.; Baric, R. S., Release of severe acute respiratory syndrome coronavirus nuclear import block enhances host transcription in human lung cells. J Virol 2013, 87, (7), 3885–902.

28. Kamitani, W.; Narayanan, K.; Huang, C.; Lokugamage, K.; Ikegami, T.; Ito, N.; Kubo, H.; Makino, S., Severe acute respiratory syndrome coronavirus nsp1 protein suppresses host gene expression by promoting host mRNA degradation. Proc Natl Acad Sci U S A 2006, 103, (34), 12885–90.

29. Bochkov, Y. A.; Hanson, K. M.; Keles, S.; Brockman-Schneider, R. A.; Jarjour, N. N.; Gern, J. E., Rhinovirus-induced modulation of gene expression in bronchial epithelial cells from subjects with asthma. Mucosal immunology 2010, 3, (1), 69–80.

30. Bosco, A.; Wiehler, S.; Proud, D., Interferon regulatory factor 7 regulates airway epithelial cell responses to human rhinovirus infection. BMC genomics 2016, 17, 76.

31. Kim, T. K.; Bheda-Malge, A.; Lin, Y.; Sreekrishna, K.; Adams, R.; Robinson, M. K.; Bascom, C. C.; Tiesman, J. P.; Isfort, R. J.; Gelinas, R., A systems approach to understanding human rhinovirus and influenza virus infection. Virology 2015, 486, 146–57.

32. Proud, D.; Turner, R. B.; Winther, B.; Wiehler, S.; Tiesman, J. P.; Reichling, T. D.; Juhlin, K. D.; Fulmer, A. W.; Ho, B. Y.; Walanski, A. A.; Poore, C. L.; Mizoguchi, H.; Jump, L.; Moore, M. L.; Zukowski, C. K.; Clymer, J. W., Gene expression profiles during in vivo human rhinovirus infection: insights into the host response. Am J Respir Crit Care Med 2008, 178, (9), 962–8.

33. Ioannidis, I.; McNally, B.; Willette, M.; Peeples, M. E.; Chaussabel, D.; Durbin, J. E.; Ramilo, O.; Mejias, A.; Flano, E., Plasticity and virus specificity of the airway epithelial cell immune response during respiratory virus infection. J Virol 2012, 86, (10), 5422–36.

34. Menachery, V. D.; Eisfeld, A. J.; Schafer, A.; Josset, L.; Sims, A. C.; Proll, S.; Fan, S.; Li, C.; Neumann, G.; Tilton, S. C.; Chang, J.; Gralinski, L. E.; Long, C.; Green, R.; Williams, C. M.; Weiss, J.; Matzke, M. M.; Webb-Robertson, B. J.; Schepmoes, A. A.; Shukla, A. K.; Metz, T. O.; Smith, R. D.; Waters, K. M.; Katze, M. G.; Kawaoka, Y.; Baric, R. S., Pathogenic influenza viruses and coronaviruses utilize similar and contrasting approaches to control interferon-stimulated gene responses. MBio 2014, 5, (3), e01174–14.

35. Wang, J.; Nikrad, M. P.; Phang, T.; Gao, B.; Alford, T.; Ito, Y.; Edeen, K.; Travanty, E. A.; Kosmider, B.; Hartshorn, K.; Mason, R. J., Innate immune response to influenza A virus in differentiated human alveolar type II cells. Am JRespir Cell Mol Biol 2011, 45, (3), 582–91.

36. Lee, S. M.; Chan, R. W.; Gardy, J. L.; Lo, C. K.; Sihoe, A. D.; Kang, S. S.; Cheung, T. K.; Guan, Y. I.; Chan, M. C.; Hancock, R. E.; Peiris, M. J., Systems-level comparison of host responses induced by pandemic and seasonal influenza A H1N1 viruses in primary human type I-like alveolar epithelial cells in vitro. Respir Res 2010, 11, 147.

37. Stegemann-Koniszewski, S.; Jeron, A.; Gereke, M.; Geffers, R.; Kroger, A.; Gunzer, M.; Bruder, D., Alveolar Type II Epithelial Cells Contribute to the Anti-Influenza A Virus Response in the Lung by Integrating Pathogen- and Microenvironment-Derived Signals. MBio 2016, 7, (3).

38. Christiansen, S. C.; Eddleston, J.; Bengtson, S. H.; Jenkins, G. R.; Sarnoff, R. B.; Turner, R. B.; Gwaltney, J. M., Jr.; Zuraw, B. L., Experimental rhinovirus infection increases human tissue kallikrein activation in allergic subjects. Int Arch Allergy Immunol 2008, 147, (4), 299–304.

39. Naclerio, R. M.; Proud, D.; Lichtenstein, L. M.; Kagey-Sobotka, A.; Hendley, J. O.; Sorrentino, J.; Gwaltney, J. M., Kinins are generated during experimental rhinovirus colds. J Infect Dis 1988, 157, (1), 133–42.

40. Hewson, C. A.; Haas, J. J.; Bartlett, N. W.; Message, S. D.; Laza-Stanca, V.; Kebadze, T.; Caramori, G.; Zhu, J.; Edbrooke, M. R.; Stanciu, L. A.; Kon, O. M.; Papi, A.; Jeffery, P. K.; Edwards, M. R.; Johnston, S. L., Rhinovirus induces MUC5AC in a human infection model and in vitro via NF-kappaB and EGFR pathways. Eur Respir J 2010, 36, (6), 1425–35.

41. Khanolkar, A.; Hartwig, S. M.; Haag, B. A.; Meyerholz, D. K.; Epping, L. L.; Haring, J. S.; Varga, S. M.; Harty, J. T., Protective and pathologic roles of the immune response to mouse hepatitis virus type 1: implications for severe acute respiratory syndrome. J Virol 2009, 83, (18), 9258–72.

42. Khanolkar, A.; Fulton, R. B.; Epping, L. L.; Pham, N. L.; Tifrea, D.; Varga, S. M.; Harty, J. T., T cell epitope specificity and pathogenesis of mouse hepatitis virus-1-induced disease in susceptible and resistant hosts. J Immunol 2010, 185, (2), 1132–41.

43. van Pesch, V.; Lanaya, H.; Renauld, J. C.; Michiels, T., Characterization of the murine alpha interferon gene family. J Virol 2004, 78, (15), 8219–28.

44. Maniatis, T.; Goodbourn, S.; Fischer, J. A., Regulation of inducible and tissue-specific gene expression. Science 1987, 236, (4806), 1237–45.

45. Marie, I.; Durbin, J. E.; Levy, D. E., Differential viral induction of distinct interferon-alpha genes by positive feedback through interferon regulatory factor-7. The EMBO journal 1998, 17, (22), 6660–9.

46. de Weerd, N. A.; Nguyen, T., The interferons and their receptors-distribution and regulation. Immunology and cell biology 2012, 90, (5), 483–91.

47. Le Goffic, R.; Pothlichet, J.; Vitour, D.; Fujita, T.; Meurs, E.; Chignard, M.; Si-Tahar, M., Cutting Edge: Influenza A virus activates TLR3-dependent inflammatory and RIG-I-dependent antiviral responses in human lung epithelial cells. J Immunol 2007, 178, (6), 3368–72.

48. Wang, Q.; Nagarkar, D. R.; Bowman, E. R.; Schneider, D.; Gosangi, B.; Lei, J.; Zhao, Y.; McHenry, C. L.; Burgens, R. V.; Miller, D. J.; Sajjan, U.; Hershenson, M. B., Role of double-stranded RNA pattern recognition receptors in rhinovirus-induced airway epithelial cell responses. J Immunol 2009, 183, (11), 6989–97.

49. Zaritsky, L. A.; Bedsaul, J. R.; Zoon, K. C., Virus Multiplicity of Infection Affects Type I Interferon Subtype Induction Profiles and Interferon-Stimulated Genes. J Virol 2015, 89, (22), 11534–48.

50. Rose, K. M.; Weiss, S. R., Murine Coronavirus Cell Type Dependent Interaction with the Type I Interferon Response. Viruses 2009, 1, (3), 689–712.

51. Versteeg, G. A.; Bredenbeek, P. J.; van den Worm, S. H.; Spaan, W. J., Group 2 coronaviruses prevent immediate early interferon induction by protection of viral RNA from host cell recognition. Virology 2007, 361, (1), 18–26.

52. Zhou, H.; Perlman, S., Mouse hepatitis virus does not induce Beta interferon synthesis and does not inhibit its induction by double-stranded RNA. J Virol 2007, 81, (2), 568–74.

53. Baig, E.; Fish, E. N., Distinct signature type I interferon responses are determined by the infecting virus and the target cell. Antivir Ther 2008, 13, (3), 409–22.

54. Yoshikawa, T.; Hill, T. E.; Yoshikawa, N.; Popov, V. L.; Galindo, C. L.; Garner, H. R.; Peters, C. J.; Tseng, C. T., Dynamic innate immune responses of human bronchial epithelial cells to severe acute respiratory syndrome-associated coronavirus infection. PLoS One 2010, 5, (1), e8729.

55. Han, M.; Chung, Y.; Young Hong, J.; Rajput, C.; Lei, J.; Hinde, J. L.; Chen, Q.; Weng, S. P.; Bentley, J. K.; Hershenson, M. B., Toll-like receptor 2-expressing macrophages are required and sufficient for rhinovirus-induced airway inflammation. The Journal of allergy and clinical immunology 2016.

56. Miura, T. A.; Holmes, K. V., Host-pathogen interactions during coronavirus infection of primary alveolar epithelial cells. J Leukoc Biol 2009, 86, (5), 1145–1151.

57. Rzepka, J. P.; Haick, A. K.; Miura, T. A., Virus-infected alveolar epithelial cells direct neutrophil chemotaxis and inhibit their apoptosis. Am JRespir Cell Mol Biol 2012, 46, (6), 833–41.

